# Synchronicity in zebrafish locomotive circuit development mediated by electrical pacemaker interneurons

**DOI:** 10.1101/2023.08.25.554828

**Authors:** Kaleb D. Miles, Chase M. Barker, Bruce H. Appel, Caleb A. Doll

## Abstract

The earliest motor output in vertebrate animals is generated by clusters of early-born motor neurons that occupy distinct regions of the spinal cord, innervating stereotyped muscle groups. Even the simplest movements require coordinated activity across these motor pools, yet motor neurons are not directly interconnected and instead project to the periphery. Instead, emerging motor circuits might be synchronized by pacemaker interneurons. We hypothesize that pacemaker interneurons are required for synchronization of motor neuron activity throughout the spinal cord, coupling motor pools through electrical gap junctions to ultimately drive coordinated motor behavior. With functional imaging in the embryonic zebrafish spinal cord, we show that ipsilateral caudal interneurons possess periodic activity profiles prior to widespread motor circuit activity that transition to synchronized Ca^2+^ events in motor neurons throughout the spinal cord. Importantly, we also show that blockade of electrical gap junctions and ablation of pioneer pacemakers leads to desynchronization in developing motor circuits. Further, we use a genetic model of hyperactivity to gain critical insight into the consequences of errors in motor circuit formation and function, finding that Fragile X syndrome (FXS) model mutant zebrafish are hyperexcitable from the earliest phases of spontaneous behavior, show reduced sensitivity to blockade of electrical gap junctions, and have increased expression of the gap junction protein Connexin 36. Taken together, our work highlights the importance of pacemakers in the development of motor circuits and suggests that the origins of hyperactivity in neurodevelopmental disorders may be established during the initiation of motor circuit formation.

## Introduction

Vertebrate locomotion in mature animals involves a complex interplay of synaptic messages, integration with sensory networks, and guidance from the brain^1–4^. Although embryonic motor circuits are far less complex, development of the motor network has been difficult to monitor and manipulate, given spontaneous motor behavior initiates in early embryonic stages^5–9^. Many nascent circuits in the CNS are influenced by central pattern generators or pacemakers that facilitate synchronization of young neuronal networks^10–12^. Electrical synapses also contribute to early locomotive networks, as gap junction blockade can completely suppress MN firing and behavioral output^8^ and prevent periodic MN activity even in the absence of interneuron pacemakers^13^. It is critical to reveal the mechanisms that synchronize early motor neuron activity to lay the groundwork for more refined behavior, as errors in motor circuit development could have persistent influence.

What are the consequences of abnormalities in motor circuit formation? One possibility is that changes in motor pacemaker function and connectivity could drive hyperactivity. Individuals with neurodevelopmental disorders, including Fragile X syndrome (FXS) and associated autism spectrum disorders (ASD), often display hyperexcitable behaviors, including repetitive movements and epilepsy^14,15^. FXS and ASD pathology appears rooted in errors in synapse formation and refinement, as shown in a variety of models and unique regions of the nervous system^16–19^. Although many therapeutics target these motor symptoms in FXS and ASD^20,21^, we still lack insight into the origins of dysfunctional motor behavior.

The zebrafish embryonic spinal cord contains a well-defined motor circuit, composed of interneurons (INs) that modulate motor neuron (MN) output to peripheral muscles. Importantly, coordinated motor output requires synchronous output from pools of MNs along the length of the spinal cord^22–24^. Previous work showed that ipsilateral caudal (IC) and ventral lateral descending (VeLD) interneurons display rhythmic activity in embryogenesis and are highly coupled to other early-born motor circuit neurons through electrical coupling^25,26^. Our work combines behavioral and targeted functional imaging approaches to reveal greater insight into locomotive circuit formation, complementing behavioral assays with foundational details at the cellular and circuit levels. We uncover additional insight on how electrical coupling mediates motor circuit communication and early behavior, showing that gap junctions facilitate coordinated locomotive output. Using a cell ablation strategy, we further demonstrate that IC pacemaker INs drive motor circuit synchronization, illustrating the critical developmental influence of pacemaker output. To investigate the basis of hyperactivity in disorders such as FXS, we also tested behavior and circuit function in *fmr1* mutant embryos. We found that *fmr1* embryos are hyperexcitable from the onset of spontaneous behavior, display increased motor circuit activity, and express surplus gap junction proteins. Taken together, our work reveals critical roles for spinal pacemaker interneurons in motor circuit formation and presents a new framework to help understand the root causes of hyperactive motor behavior in FXS.

## RESULTS

### Initial zebrafish locomotive behavior is driven by electrical gap junction coupling

Motor behavior in zebrafish embryos initiates as spontaneous sensory-independent coiling contractions at 17.5 hours post-fertilization (hpf), when embryos are still confined to their transparent chorion membranes^27,28^. Importantly, these early contractions are not dependent on chemical neurotransmission and are instead rooted in electric coupling through gap junctions, as movement ceases completely upon application of concentrated heptanol (2 mM), a gap junction blocker^8^, presumably through a complete blockade of gap junctions. To gain more insight into the subtleties of electrical coupling on motor circuit output, we applied subthreshold heptanol and scored embryonic behavior, finding that 50 μM heptanol exposure did not prevent locomotive output. Instead, this concentration drove a prominent increase in “weak” uncoordinated coiling movements (Figure 1A,B,D, Videos 1,2), and overall hyperactivity due to a prominent increase in these weak movements (Figure 1E). This suggests that gap junction coupling facilitates coordinated activity and partial blockade can spur uncoordinated hyperactivity. Weak coiling behavior was even more prominent with application of 100 μM heptanol, but this concentration greatly reduced overall activity (Figure 1B,C,E, Video 3). This data complements previous physiological work showing electrical coupling regulates early motor circuit activity^8,25^, and also adds an important role for gap junctions in promoting coordinated locomotive output.

**Figure 1.**
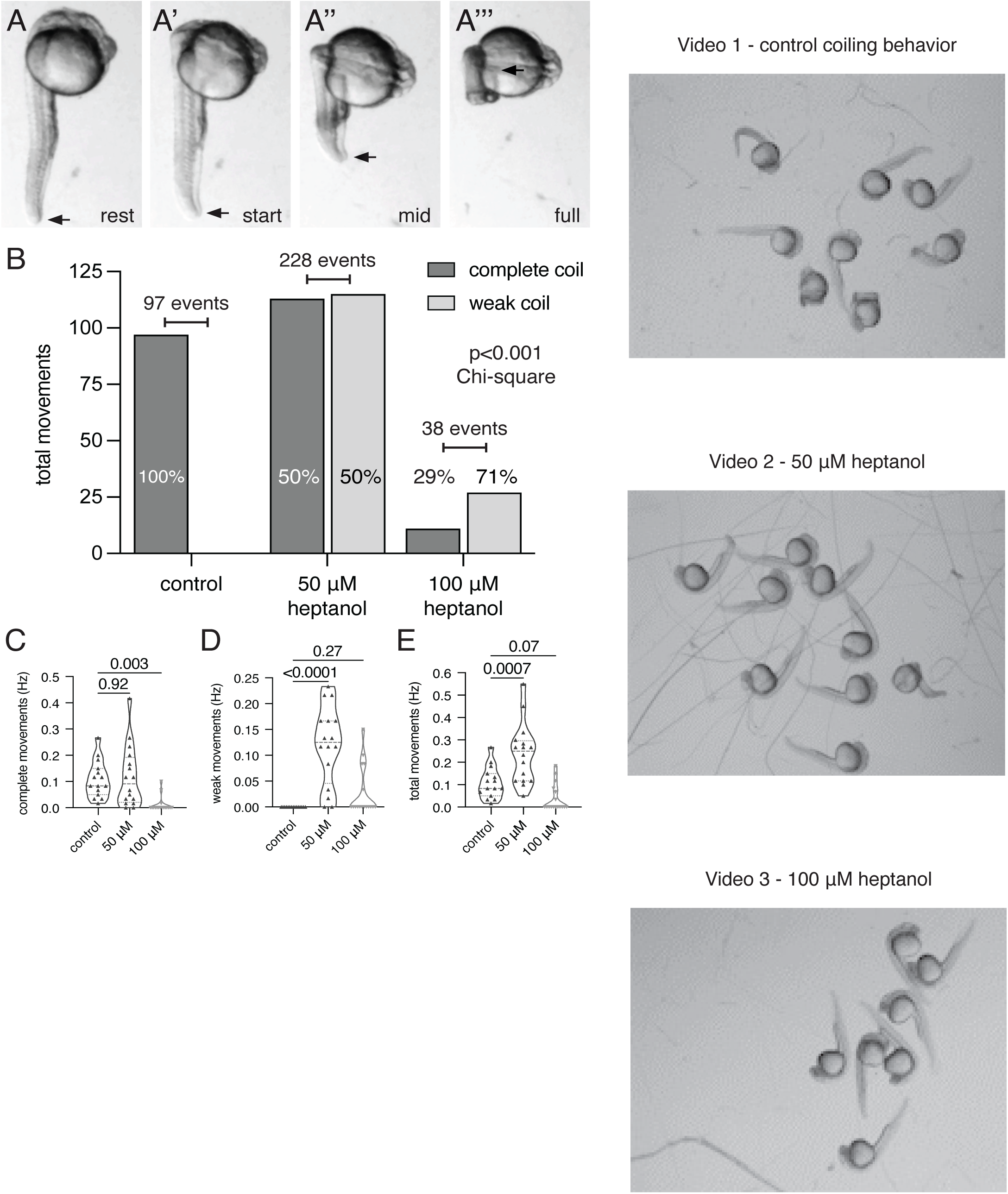
Gap junctions mediate spontaneous embryonic coiling behavior. Representative still frames images illustrate embryonic coiling behavior, in which the embryo tail undergoes a progressive movement from rest (A), coil initiation (A’), mid-coil (A’’), to complete coil at the level of the hindbrain (A’’’). (B) Contingency plot displaying the effects of increased heptanol exposure on embryonic coiling behavior, comparing weak and complete coils in control embryos, and embryos exposed to 50 and 100 μM heptanol, an electrical gap junction blocker (Chi-square test of proportions; control=97 total events, [50]=228, [100]=38). Graphs show complete movement frequency (C; control=0.11±0.02 Hz; [50]=0.12±0.03 Hz; [100]=0.01±0.01 Hz), weak movement frequency (D; control=0±0 Hz; [50]=0.12±0.02 Hz; [100]=0.03±0.01 Hz), and cumulative movement frequency (E; control=0.11±0.02 Hz; [50]=0.24±0.03 Hz; [100]=0.04±0.02 Hz; significance by ANOVA tests with Dunnett’s multiple comparison tests. nControl=15 embryos; n[50]=16; n[100]=17. See Videos 1-3.

### Spinal interneurons are highly coupled in the early locomotive network

How do disparate pools of motor neurons (MNs) organize to generate coordinated behavior? Remarkably, spinalization has no effect on early spontaneous zebrafish activity, indicating that early motor circuits are not regulated by the brain and instead rely on local spinal networks^8,27,29^. The cellular framework underlying the initiation of motor behavior in zebrafish in driven by three cardinal types of primary MNs in zebrafish that undergo axogenesis into unique muscle groups to drive early motor behavior^9,24,30,31^. In addition, two interneuron (IN) classes contribute to embryonic motor activity: ipsilateral caudally projecting (IC) cells are positioned in the caudal hindbrain/rostral spinal cord with limited axonal extensions into rostral somites^32^, and ventrolateral descending (VeLD) INs, which are highly similar in morphology and physiology to ICs and are born alongside MNs throughout the spinal cord^25,33^. As the gene *mnx1* is specifically expressed in motor circuit neurons, we used *mnx1* transgenic tools to examine cellular organization and relationships. At the stage corresponding to initial spontaneous behavior (∼18 hpf), pools of *mnx1^+^* cells were distributed in defined hemisegments of the spinal cord, though very few MNs had begun axogenesis this stage (Figure 2A,B, see also Figure 6A,B)^9^. In contrast, IC cell axons pioneer the medial longitudinal fasciculus (MLF)^32^, and in transgenic embryos expressing the calcium (Ca^2+^) sensor GCaMP-HS in *mnx1^+^*cells (*mnx1:*GAL4;*UAS:*GCaMP-HS), IC axons were apparent in the ventral spinal cord, passing near the somata of caudal *mnx1+* cells (Figure 2B)^34^. Therefore, IC neurons are morphologically positioned to influence early born motor circuit neurons.

**Figure 2.**
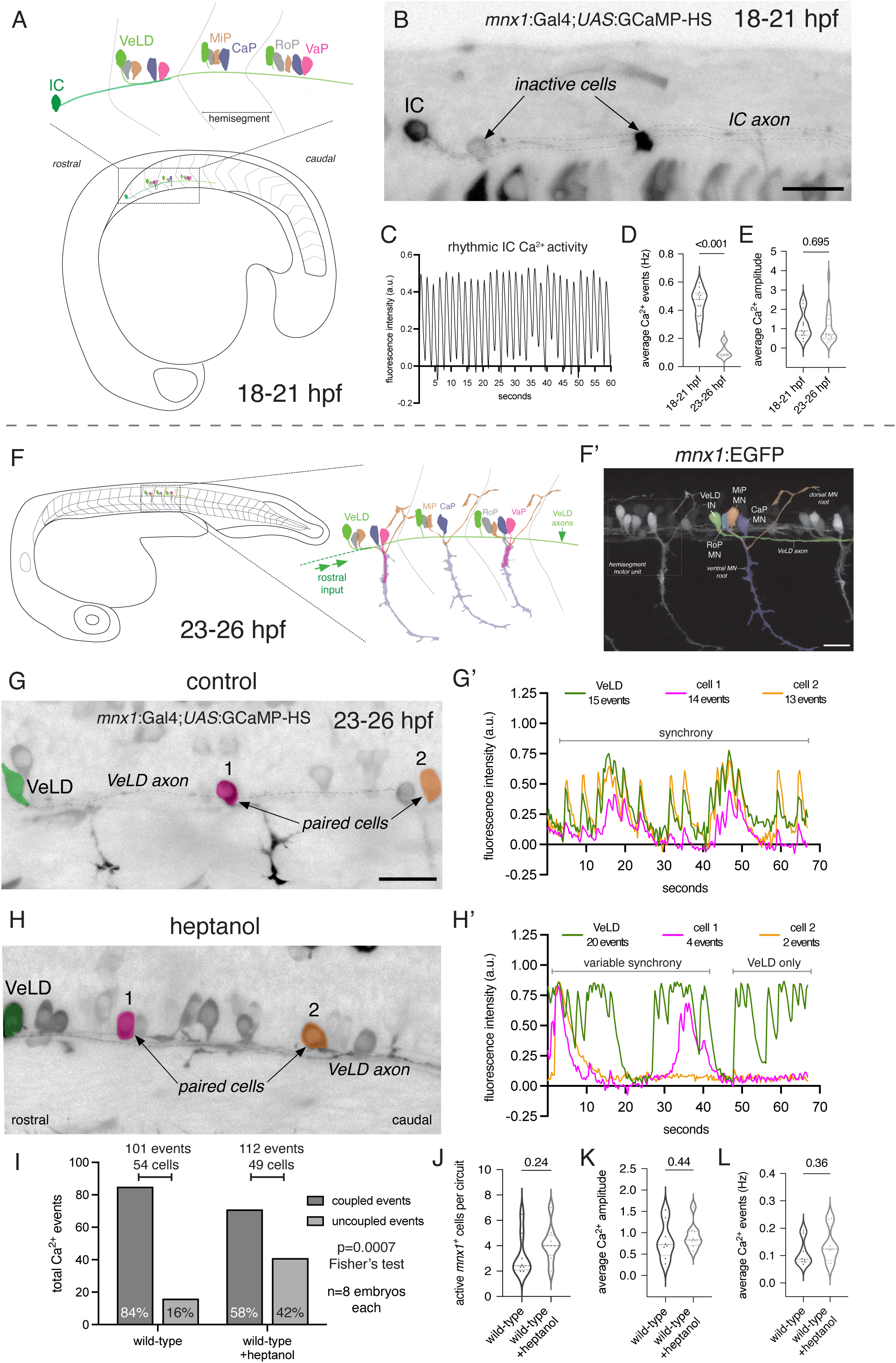
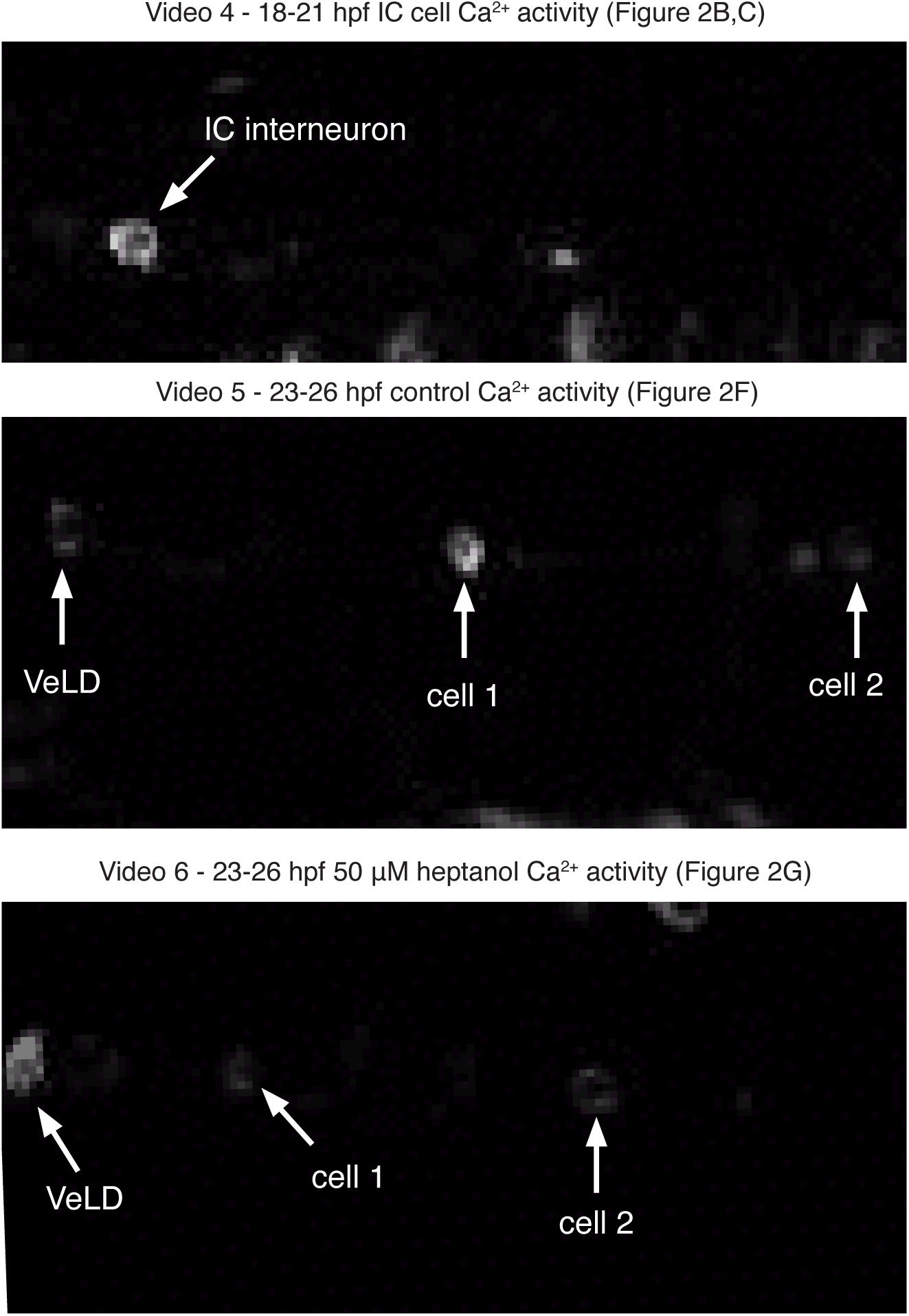
Gap junction coupling synchronizes cells in the developing zebrafish motor circuit. (A) Schematic showing motor circuit organization in early embryogenesis (18-21 hpf): ipsilateral caudal interneurons (IC; dark green) in the caudal hindbrain/rostral spinal cord send axon projections caudally to rostral spinal cord hemisegments, which contain motor pools of primary motor neurons and ventral lateral descending interneurons (VeLD, light green). (B) Static image of *mnx1:*GAL4;*UAS:*GCaMP-HS+ cells in the spinal cord at the 18-21 hpf stage, including an IC cell displaying rhythmic Ca2+ activity (see 2C) and ipsilateral silent neurons (image color inverted to highlight neuronal projections). (C) Chart showing rhythmic Ca2+ transients in an IC cell at the 18-21 hpf stage. Graphs displaying the average number of Ca2+ event frequency (D; 18-21 hpf=0.45±0.03 Hz; 23-26 hpf=0.11±0.02 Hz) and average peak Ca2+ amplitude (E; 18-21 hpf=1.19±0.19 arbitrary units, n=13 embryos, only IC cells; 23-26 hpf=1.07±0.19 a.u., n=20 embryos, *mnx1*+ cells). (F) Schematic showing motor circuit organization at the 23-26 hpf stage: each hemisegment of the spinal cord contains primary motor neurons (rostral primary, RoP; middle primary, RoP; caudal primary, CaP; variable primary, VaP) that innervate specific muscle groups. VeLD INs have extensive axons that pass through many caudal hemisegments. (F’) *mnx1*:EGFP transgenic embryo at 24 hpf, pseudocolored to highlight motor circuit neurons. (G) Still frame image of a *mnx1:*GAL4;*UAS:*GCaMP-HS transgenic motor circuit at the 23-26 hpf stage, showing an active VeLD IN (green; VeLD axon indicated by dashed line) and two caudal ipsilateral cells (pink, orange). (G’) Corresponding functional imaging plot shows tightly linked Ca2+ activity in VeLD INs and ipsilateral cells. (H) Image of a transgenic motor circuit following 1-hour 50 μM heptanol exposure, including an active VeLD and two ipsilateral cells. (H’) Corresponding graph shows that heptanol exposure desynchronized VeLD activity from ipsilateral cells, as not all VeLD Ca2+ events were matched in caudal neurons. Quantification of the impact of heptanol exposure on motor circuit synchronization, including the relative proportion of paired and unpaired transients (I), average active *mnx1+* cells per circuit (J; control=3.21±0.58 cells; heptanol=4.17±0.52 cells), average Ca2+ event amplitude (K; control=0.83±0.15 arbitrary units; heptanol=0.92±0.11 a.u.), and average Ca2+ event frequency (L; control=0.11±0.02 Hz; heptanol=0.14±0.02 Hz). n=8 embryos each condition (I-L). Scale bars: 10 μm (A), 20 μm (B, F, G). See Videos 4-6.

We used functional Ca^2+^ imaging to examine activity profiles in *mnx1^+^* motor circuit cells in early embryogenesis and to visualize the influence of IN pacemakers on locomotive circuit output. Prior work has shown that IC and VeLD physiology fits the description of pacemakers, as these INs display Ca^2+^- dependent rhythmic depolarizations and are electrically coupled with ipsilateral MNs and VeLDs^25,26^. An additional study showed that motor circuit neurons display synchronized activity as spontaneous coiling initiates^35^, though this work was unable to distinguish IN activity from other *olig2*-expressing MNs. At the 18-21 hpf stage, we detected very few Ca^2+^ events, other than a bilateral pair of *mnx1^+^* IC cells in the rostral spinal cord, which showed robust periodic activity profiles, pulsing at an average rate of ∼0.45 Hz with average Ca^2+^ event amplitude of ∼1.2 arbitrary units (Figure 2C,D,E; Video 4)^25,26^. Although GCaMP-HS expression was apparent in cells ipsilateral and caudal to active IC cells at this stage (presumptive VeLDs/MNs), very few of these cells showed Ca^2+^ activity, especially at the early portion of this developmental window. This data supports the hypothesis that ICs function as motor pacemakers.

More refined locomotive output would require additional synchronized activity from MN pools throughout the spinal cord. Embryonic motor activity transitions from spastic movements into complete body coils between 18 and 24 hpf^27,28^ (see also Videos 9-12), and MN axogenesis is dramatic during this transition, as at 24 hpf the unique *mnx1^+^* neuronal subtypes had characteristic axonal arbors and were stereotypically arranged at the level of the mid-trunk (Figure 2F)^36,37^. Integrative pacemaker interneurons would also be required at this region to synchronize distinct pools of MNs, yet IC IN axons only extend through ∼10 somites at the rostral end of the spinal cord ^25^. Instead, *mnx1^+^* VeLD INs were evident in the mid trunk motor circuit, with axons that traversed caudally through the medial longitudinal fasciculus (MLF). VeLD axons passed through many successive motor pools near MN cell bodies and also overlapped with additional ipsilateral VeLDs (Figure 2F; green pseudocolor)^25,33,36^. Given commonalities in morphology and physiology with IC cells, VeLDs could represent secondary pacemakers^25,26^. Also fitting the transition to more coordinated motor output, Ca^2+^ signaling dynamics changed dramatically over the course of just a few hours, with far more active *mnx1^+^* neurons at ∼23 hpf at the level of the mid-trunk. Overall, *mnx1*^+^ cells displayed reduced periodicity at this later stage, with fewer average Ca^2+^ events (∼8.1 Hz; Figure 2D; in line with prior work ^26^), but no change in Ca^2+^ event amplitude over development (Figure 2E). This data shows that motor circuit activity patterns change quickly as locomotive output is refined, with widespread *mnx1^+^* cell activity in the spinal cord at 24 hpf.

We next tested roles for electrical gap junction coupling in developing motor circuits at the cellular level, using subthreshold heptanol exposure and Ca^2+^ imaging to complement our behavioral assays. We recorded Ca^2+^ events in *mnx1^+^* MNs and VeLD INs in the mid trunk at 23-26 hpf and determined whether Ca^2+^ events in individual cells were synchronized within their ipsilateral circuit. VeLD INs and their caudal axons were apparent in our recordings, which represent the presumptive link between *mnx1^+^* cells within the circuit (see Video 5). In control *mnx1:*GAL4;*UAS:*GCaMP-HS embryos, 84% of Ca^2+^ events were synchronized with another *mnx1^+^* cell in the ipsilateral circuit (Figure 2G; Video 5), indicating neuronal activity was tightly linked in motor circuit cells. In contrast, only 63% of Ca^2+^ events were synchronized following 1-hour exposure of embryos to 50 μM heptanol (Figure 2H; Video 6). This represents a prominent desynchronization of *mnx1^+^* Ca^2+^ events, in an effect that was comparable to broad optogenetic silencing of motor circuit neurons in a prior study ^35^. Heptanol exposure had no effect on the average number of active *mnx1^+^* cells per circuit, Ca^2+^ event amplitude, or the number of Ca^2+^ events per cell (Figure 2I-K), which suggests gap junction blockade may isolate neurons from the IN pacemaker without influencing neuronal physiology. Taken together with behavioral data, this suggests that gap junctions mediate motor circuit synchronization with pacemakers to achieve coordinated motor output.

### Early pacemaker activity guides subsequent motor circuit synchronization

IC interneurons appear to represent pioneer pacemakers that entrain activity in the rest of the spinal cord, as they possess physiological profiles that fit pacemaker identity^25,26^ and showed the earliest activity in our imaging paradigm (Figure 2). To gain greater cell and circuit-level insight on the influence of pioneer pacemakers, we laser ablated IC cells prior to widespread circuit activity and then gauged subsequent motor circuit synchronization. Due to the ambiguity of early-born *mnx1*^+^ cell identity prior to axogenesis, we performed these experiments in transgenic *mnx1:*GAL4*;UAS:*GCaMP-HS embryos, focusing on the bilateral IC cells in the rostral spinal cord displaying periodic Ca^2+^ activity at 18-20 hpf (Figure 3A,B). To maximize circuit data collection, we mounted embryos dorsally and then selectively removed one IC from each side of the embryo with targeted laser ablation (Figure 3C)^38^. Embryos were allowed to recover for ∼4 hours (23-25 hpf), then remounted to record widespread Ca^2+^ activity in *mnx1^+^*cells in ipsilateral motor circuits on each half of the spinal cord.

**Figure 3.**
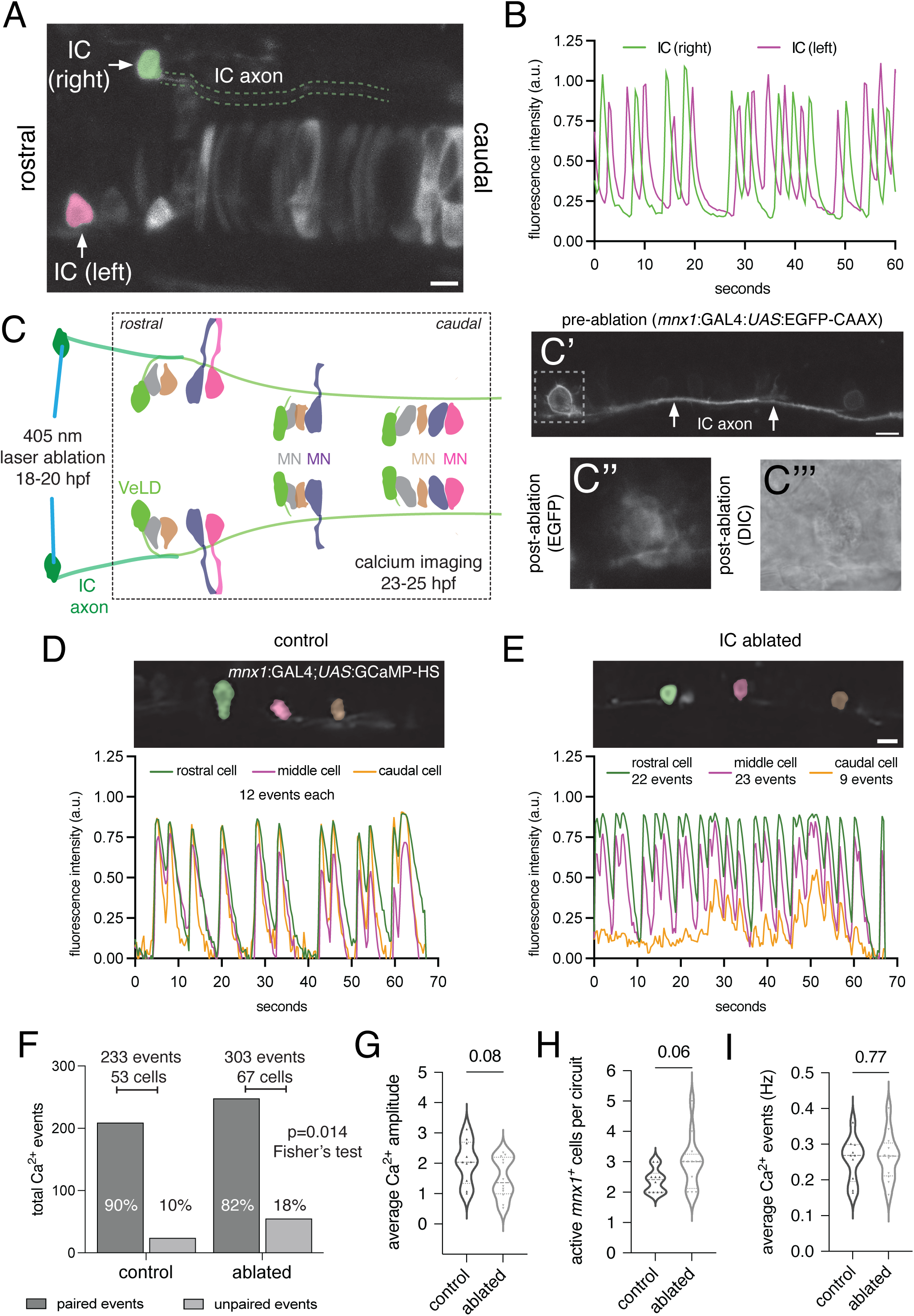
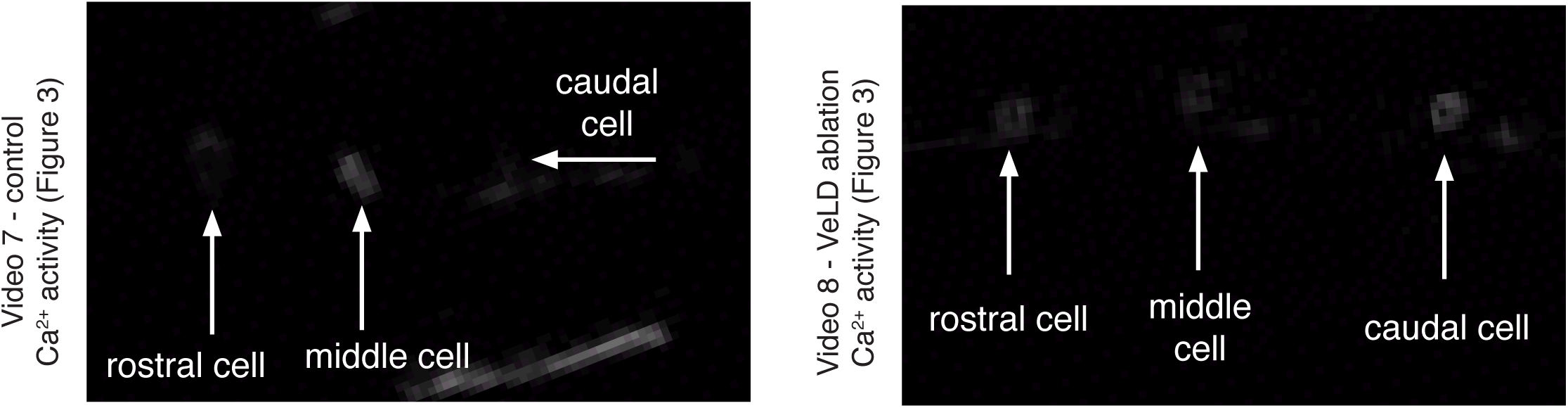
Removal of pacemaker interneurons disrupts motor circuit synchronization. (A) Representative still frame image from the dorsal perspective showing active *mnx1:*GAL4;*UAS:*GCaMP-HS+ ipsilateral caudal (IC) INs in both the right (green) and left (magenta) sides of the rostral spinal cord (a portion of the caudal IC axon is highlighted by dashed lines). (B) Corresponding charts show alternating fluorescence intensity changes in GCaMP+ IC INs on each half of the spinal cord. (C) Experimental schematic from the dorsal perspective (as in A), where an active GCaMP-HS+ IC IN is selectively laser ablated at 18-20 hpf on each half of the spinal cord, followed by capture of circuit level Ca2+ dynamics at 23-25 hpf. Representative proof of principle images in an embryo expressing *mnx1:*GAL4;*UAS*:EGFP-CAAX show an IC cell prior to ablation (C’) and zoomed images after 45 seconds of 405 nm laser exposure (C’’, EGFP; C’’’, DIC). (D) Example control circuit with three synchronized motor circuit neurons. (E) Example motor circuit following pacemaker IC cell ablation, in which the caudal cell (orange) is desynchronized from rostral cells (green, pink). (F) Contingency plot showing the influence of early IC cell ablation on motor circuit synchronization at 23-25 hpf (percentages in bars indicate the percentage of total events in each condition). Quantification of average Ca2+ amplitude (G; control=2.04±0.22 arbitrary units; ablated=1.50±0.19 a.u.), active cells per circuit (H; control=2.38±0.12 cells; ablated=2.99±0.25 cells), and Ca2+ event frequency (I; control=0.26±0.02 Hz; ablated=0.27±0.02 Hz). nControl=10 embryos, nAblated=12 (F-I). IC=ipsilateral caudal interneuron; VeLD=ventral lateral descending interneuron; MN=motor neuron. Scale bars = 10 μm. Statistical significance determined by t-tests, where p<0.05 is significant.

We found that removal of early-active IC INs led to subsequent desynchronization in the cord. While most Ca^2+^ events in controls were paired with other *mnx1^+^* cells in their circuit (Figure 3D,F), early IC ablation led to an increase in uncoupled Ca^2+^ activity following ablation compared to controls (Figure 3E,F; Videos 7,8), similar to the effects of heptanol decoupling (Figure 2). Early IC removal also led to a slight reduction in *mnx1^+^* cell Ca^2+^ amplitude and an increase in active cells per circuit, though these effects did not reach significance (Figure 3G,H). It is important to note that persistent activity following early IC removal suggests considerable redundancy in the developing motor circuit, which may come from secondary VeLDs pacemakers. However, these targeted experiments show that pioneer IC interneurons perform a critical developmental role in motor circuit synchronization.

### fmr1 mutant embryos display hyperexcitable behavior

What are the consequences for abnormalities in motor circuit pacemaker function? We next tested whether changes in pacemaker output or electrical connectivity contribute to a genetic model of hyperactivity. Some of the most debilitating symptoms of FXS are hyperexcitable motor behaviors^14,39,40^, which are recapitulated in a variety of models^41–43^ including the zebrafish *fmr1^hu^*^2787^ mutants, which display increased locomotion in both larval stages and in adults^44–46^. To test whether hyperactivity is also noted in embryonic stages, we assayed spontaneous embryonic behavior at 19 hpf and 24 hpf, recording 80-second videos of wild-type and *fmr1* mutant embryos still confined to their chorion membranes (Figure 4A; Videos 9-12, *in supplementary material*). Consistent with previous reports, coiling activity peaked in wild-type at 19 hpf and was less pronounced at 24 hpf. *fmr1* mutant embryos were more active than wild-type controls at both stages, with increased relative time in motion and coiling frequency (Figure 4B,C). As another indicator of embryonic motor behavior, we also quantified chorion hatching, a variable process that occurs between 48 and 72 hpf in response to the relative activity of the embryo and enzymatic digestion of the chorion^28,47^, finding a ∼29% increase in hatching in *fmr1* mutant embryos at 48 hpf (Figure 4D).

**Figure 4.**
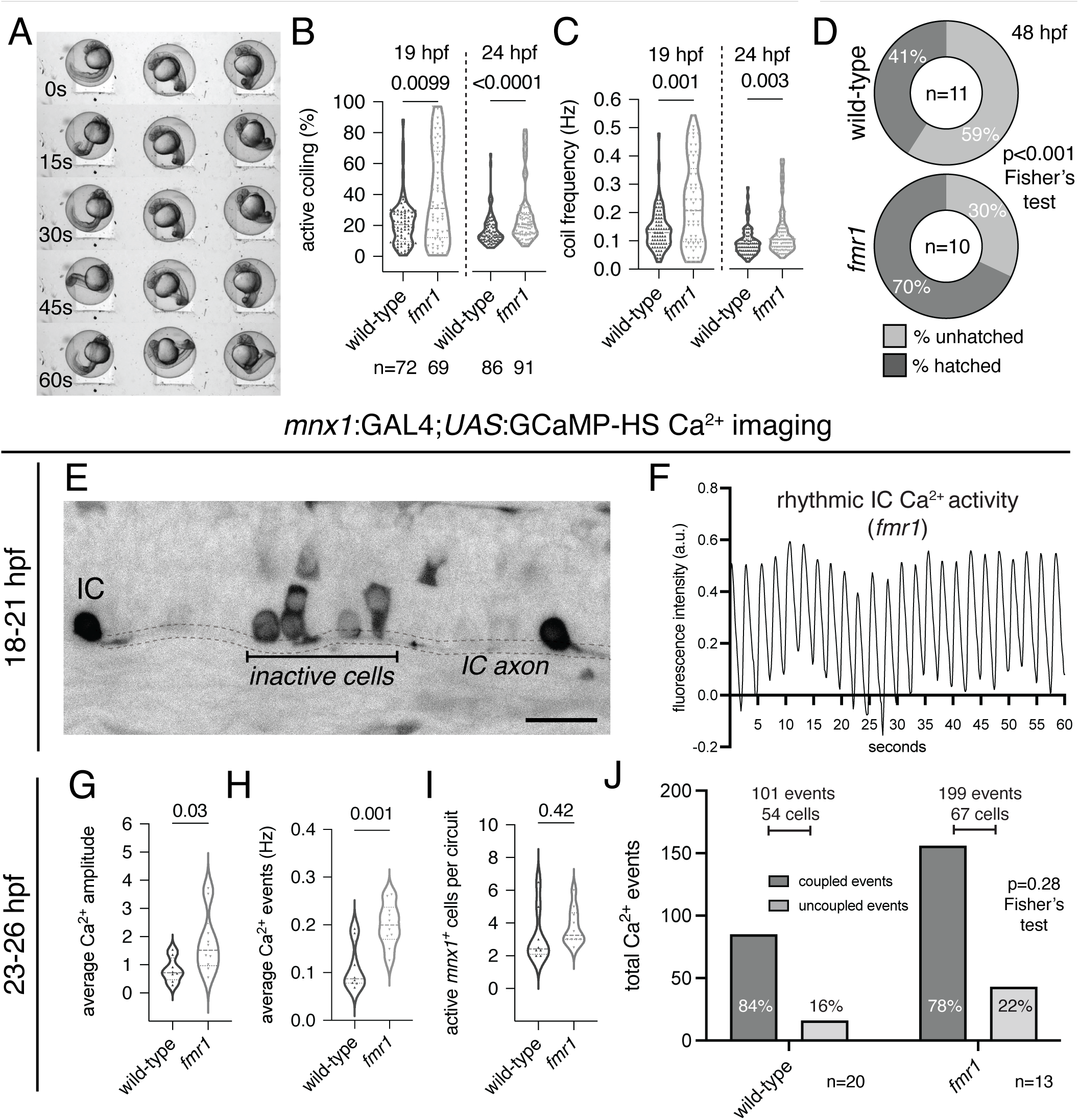
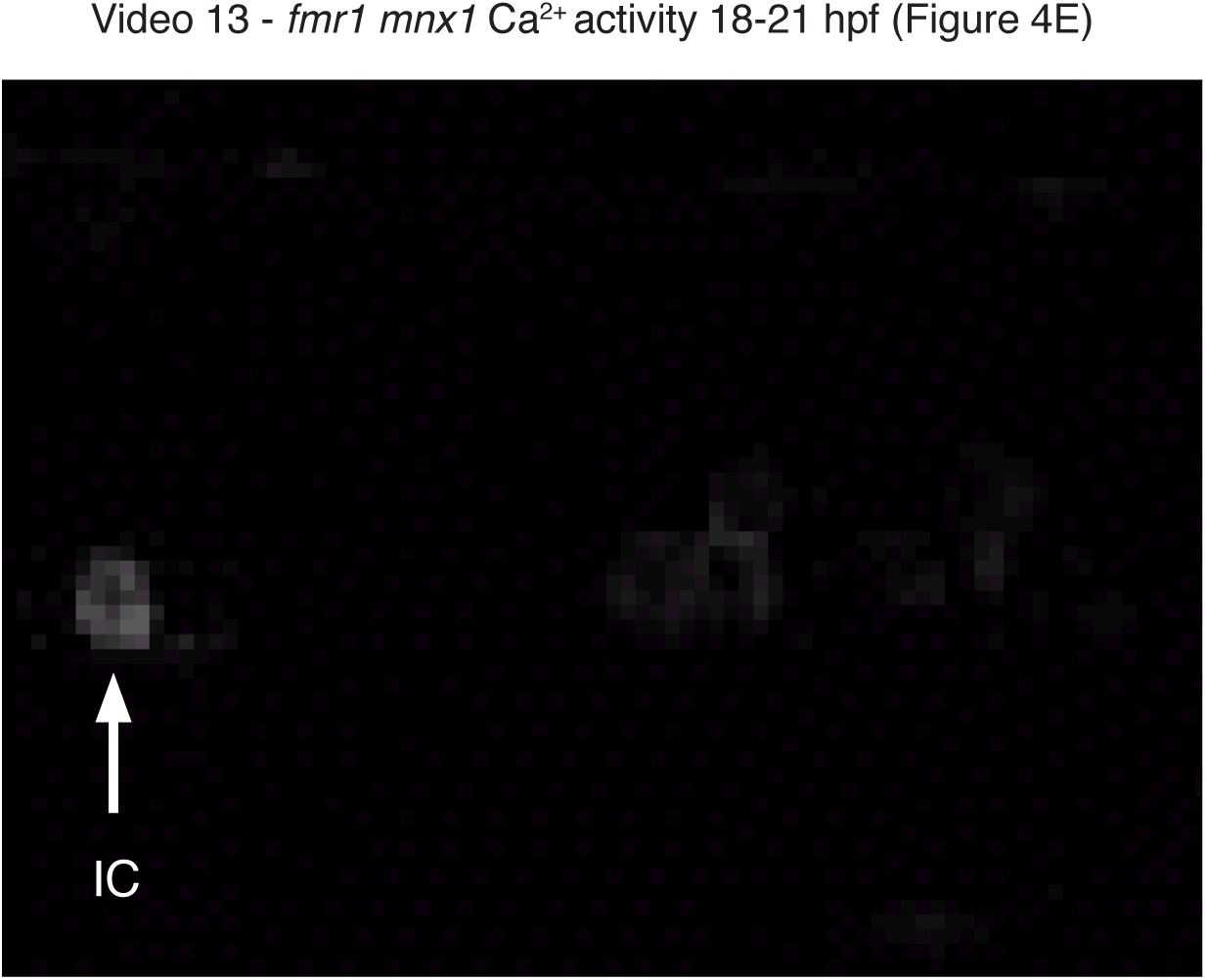
*fmr1* mutant embryos display hyperexcitable behavior and altered motor circuit activity. (A) Example still frames from embryonic behavioral videos showing spontaneous coiling movements within chorions. Quantification of embryonic behavior at 19 hpf and 24 hpf, including percentage of time in motion (B; wild-type19hpf=22.13±1.82%; *fmr1*19hpf=38.52±3.85%; wild-type24hpf=16.99±1.10%; *fmr1*24hpf=26.65±1.99%), and coiling frequency (C; wild-type19hpf=0.14±0.009 Hz; *fmr1*19hpf=0.23±0.02 Hz; wild-type24hpf=0.095±0.005 Hz; *fmr1*24hpf=0.13±0.009 Hz; significance determined by Mann-Whitney tests). (D) Quantification of chorion hatching at 48 hpf (nWT=11 clutches, 471 embryos; n*fmr1*=10 clutches, 354 embryos; significance by Fisher’s exact test of proportions). (E) Still frame image of an *fmr1* mutant embryo expressing *mnx1*:Gal4;*UAS*:GCaMP-HS at the 18-21 hpf stage, with an active IC neuron and inactive caudal ipsilateral cells. (F) Corresponding graph shows rhythmic Ca2+ periodicity in this IC cell. Quantification of average *mnx1+* Ca2+ amplitude (G; wild-type=0.83±0.15 arbitrary units; *fmr1*=1.79±0.34 a.u.), Ca2+ event frequency (H; wild-type=0.11±0.02 Hz; *fmr1*=0.20±0.01 Hz), and average number of active cells per circuit (I; wild-type=3.21±0.58 cells; *fmr1*=3.75±0.35 cells) in *fmr1* mutants compared to wild-type at 24 hpf (significance determined by t-tests). (J) Contingency plot shows the influence of heptanol exposure on motor circuit synchronization in *fmr1* mutants (significance by Fisher’s exact test of proportions). nWT=20 embryos; n*fmr1*=13 (G-J). p<0.05 is considered significant. Scale bar =10 μm.

Changes in early motor circuit activity could directly influence hyperexcitable behavior in FXS. To examine this possibility, we performed Ca^2+^ imaging in developing *fmr1* mutant motor circuits, first finding periodic Ca^2+^ activity in *fmr1* mutant IC pacemaker cells at 18-21 hpf (Figure 4E,F, Video 13). We next assayed circuit level activity in *fmr1* mutants at 23-26 hpf, finding increased Ca^2+^ amplitude and increased Ca^2+^ events per cell compared to wild-type (Figure 4G,H), indicative of hyperactive circuit output. There was no change in the average active cells per circuit (Figure 4I), nor in the relative proportion of synchronized and uncoupled *mnx1^+^* Ca^2+^ events in wild-type and *fmr1* circuits (Figure 4J). Overall, this data shows that *mnx1^+^* cells in *fmr1* mutants display hyperactive Ca^2+^ activity profiles, which is directly in line with hyperexcitable behavior noted in this FXS model.

### fmr1 mutant embryos and motor circuits are resistant to gap junction blockade

Gap junction coupling mediates motor circuit synchronization and behavioral output, and FXS model zebrafish embryos display hyperexcitability and elevated motor circuit activity. To test whether changes in electrical coupling contribute to these phenotypes, we first assayed *fmr1* mutant embryonic behavior following exposure to heptanol. Like wild-type, *fmr1* mutant embryos displayed more weak contractions in the presence of subthreshold levels of the gap junction blocker heptanol (Figure 5A,D) with no change in complete coils (Figure 5B). Importantly, this concentration did not induce hyperactivity in *fmr1* mutants, as seen in controls (Figure 5C; see Figure 1E). Although *fmr1* mutants are hyperactive at baseline (see Figure 5B,C), this suggests altered gap junction sensitivity may contribute to the hyperexcitability phenotype. Complete motor circuit electrical blockade was still possible in *fmr1* embryos, as exposure to 100 μM heptanol almost completely suppressed motor activity in *fmr1* (Figure 5A,C, Video 16), in an effect akin to wild-type (Figure 1E). Taken together, our data show that *fmr1* mutants display hyperactive behavior from the onset of spontaneous motor activity that is less sensitive to electrical gap junction blockade.

**Figure 5.**
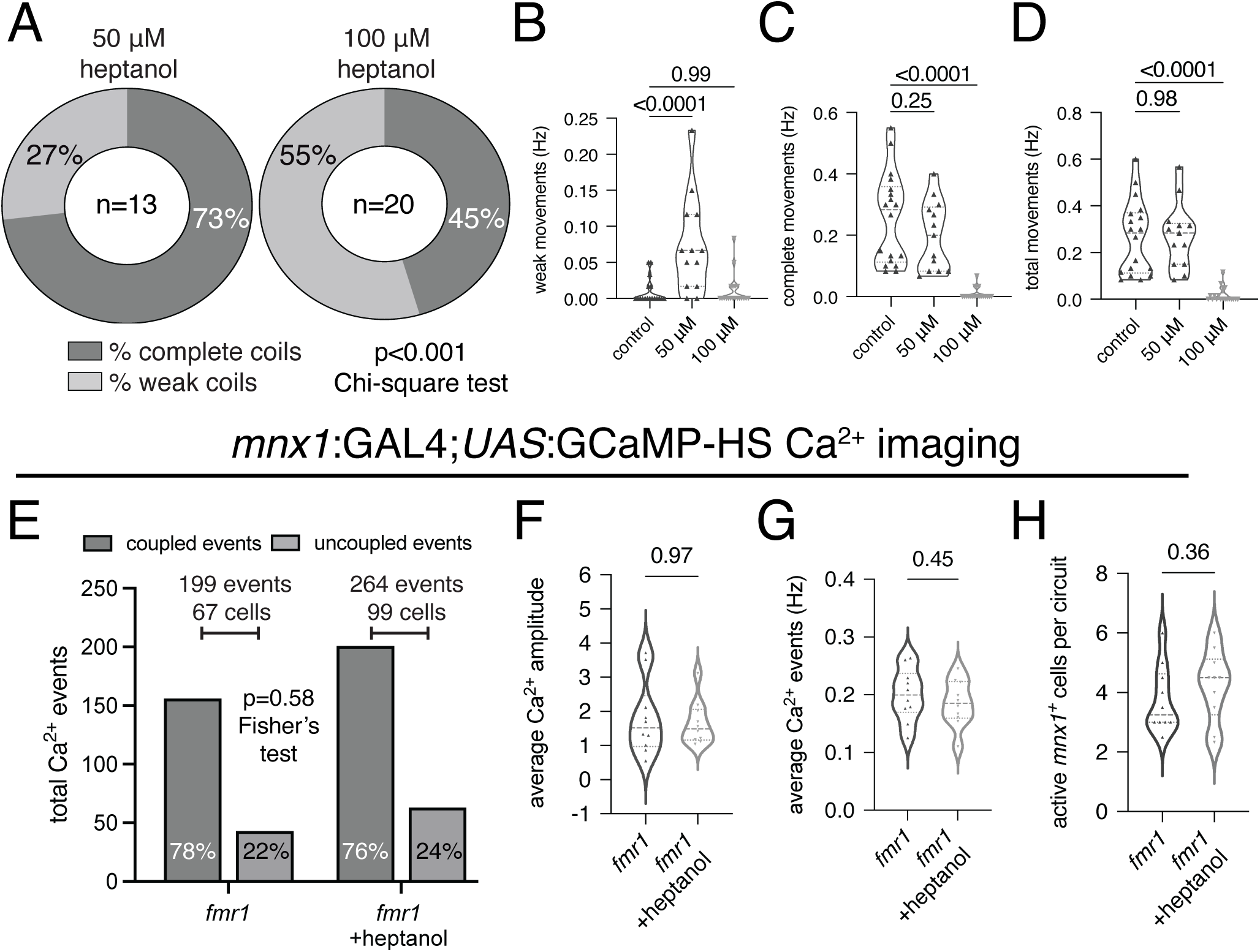
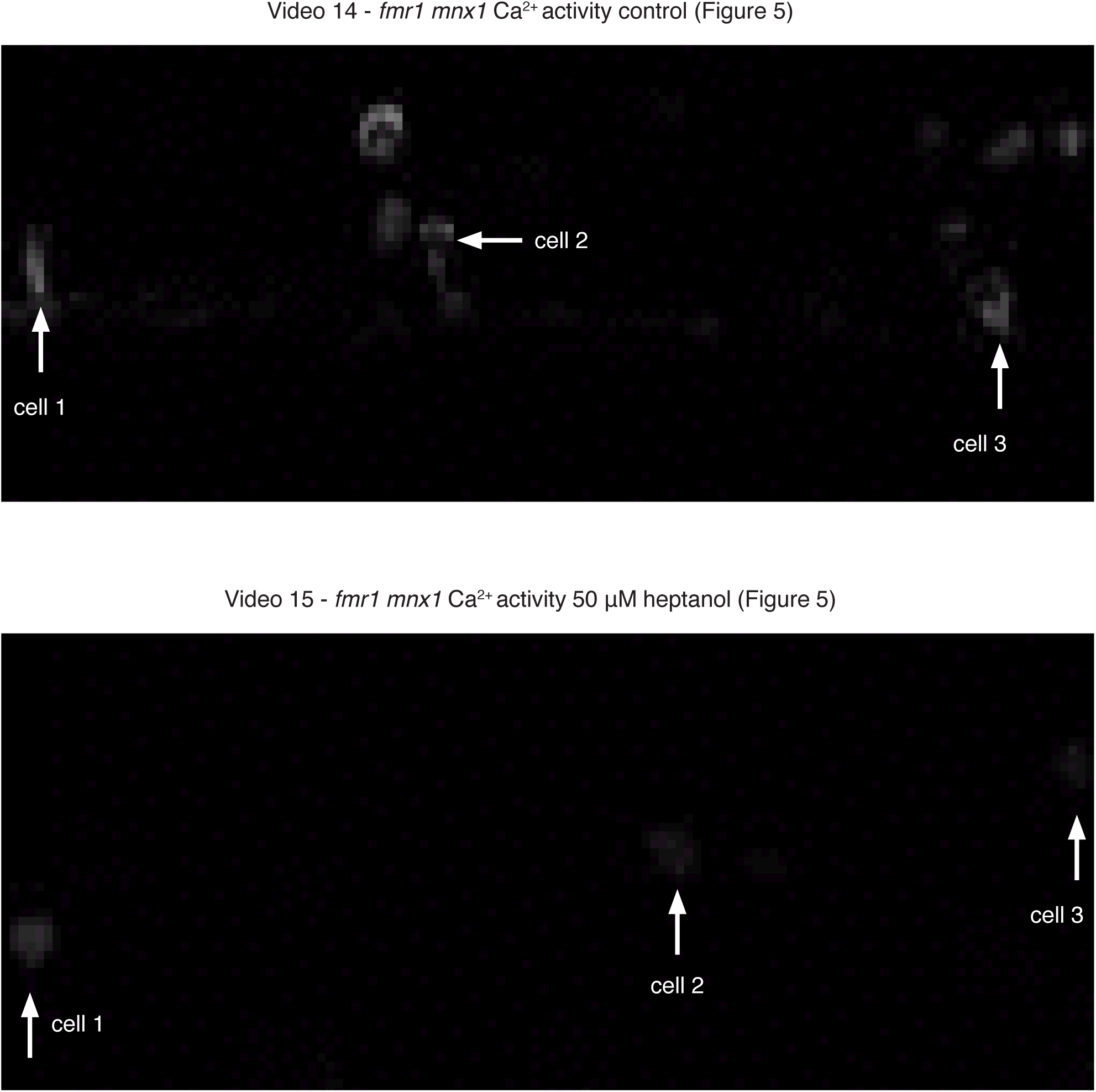
*fmr1* mutant motor circuits are resistant to gap junction blockade. (A) Behavioral effects of the electrical gap junction inhibitor heptanol on *fmr1* embryos, comparing the percentage of complete and weak coiling movements (∼97% of controls displayed complete coils; control=286 total events, [50]=115, [100]=22). Graphs compare the average number of weak movement frequency (B; control=0.009±0.004 Hz; [50]=0.07±0.02 Hz; [100]=0.01±0.005 Hz), complete movement frequency (C; control=0.26±.04 Hz; [50]=0.2±0.03 Hz; [100]=0.008±0.004 Hz), and cumulative movement frequency (D; control=0.27±0.04 Hz; [50]=0.27±0.04 Hz; [100]=0.018±0.007 Hz) in each condition (significance by ANOVA tests with Dunnett’s multiple comparison tests; nControl=18 embryos, n[50]=13, n[100]=20). (E) Contingency plot showing the influence of heptanol exposure on the relative proportion of paired and unpaired Ca2+ events in *fmr1* mutants expressing *mnx1:*GAL4;*UAS:*GCaMP-HS at 23-26 hpf (percentages in bars indicate percentage of total events; significance by Fisher’s exact test of proportions). Quantification of average Ca2+ amplitude (F; *fmr1*=1.79±0.34 arbitrary units; heptanol=1.66±0.20), Ca2+ event frequency (G; *fmr1*=0.20±0.01 Hz; heptanol=0.19±0.01 Hz), and active *mnx1+* cells per circuit (H; *fmr1*=3.75±0.35 cells; heptanol=4.23±0.38 cells) in *fmr1* mutant controls and *fmr1* mutants exposed to heptanol. n=10 embryos each (E-H). p<0.05 is considered significant. Scale bar = 10 μm.

Are changes in motor circuit heptanol sensitivity in *fmr1* mutants mirrored at the circuit level? We next tested the influence of gap junctions on synchronicity in early motor circuits in *fmr1* embryos expressing the GCaMP-HS Ca^2+^ sensor. We found that 78% of Ca^2+^ events in *mnx1^+^* cells were synchronized neurons in control *fmr1* mutants (Figure 5E, Video 14), thereby showing tight coupling with ipsilateral cells that was comparable to wild-type (Figure 2). Interestingly, 76% of Ca^2+^ events in motor circuit cells were paired in ipsilateral cells following 1-hour exposure to 50 μM heptanol (Figure 5E, Video 15), indicating that subthreshold gap junction blockade did not influence Ca^2+^ event pairing in *fmr1* mutant motor circuit cells. There was also no effect of heptanol exposure on Ca^2+^ event amplitude or *mnx1^+^* cell Ca^2+^ activity in *fmr1* mutants (Figure 5F-H). Taken together with behavioral data, these data are consistent with the possibility that electrical coupling is more robust in *fmr1* mutant embryos and may contribute to hyperexcitable behavior.

### Increased expression of Connexin 36 in the absence of Fmrp is driven by electrical coupling

One possible explanation for the altered sensitivity to heptanol in *fmr1* mutants could be a change in the density of gap junctions expressed by motor circuit neurons. To test this possibility, we used an antibody raised against the neuronal gap junction protein Connexin 36 (Cx36), which detects Connexins encoded by multiple genes in zebrafish^48,49^. We detected Cx36 expression in whole-mount wild-type and *fmr1* mutant embryos expressing the *mnx1:*EGFP transgene in these experiments to clearly label early motor units (Figure 6A,B), and used a three-dimensional surface modeling strategy to distinguish Cx36 punctal expression that was specifically localized to motor circuit neurons^50^. We first examined Cx36 expression at 17 hpf, just prior to most MN axogenesis and the initiation of spontaneous behavior, finding no change in Cx36 density on *mnx1^+^* somata in *fmr1* embryos, when normalized to the relative volume of *mnx1:*EGFP (Figure 6B,C). At 24 hpf, Cx36 expression on *mnx1*^+^ cells in the ventral spinal cord was both perisomatic and localized on MN axons (Figure 6D,E), which also form electrical synapses with muscle^51^. In wild-type, Cx36 density at 24 hpf (∼15 puncta per volume) was comparable to the 17 hpf stage (∼14 puncta per volume; Figure 6C,H), indicating no prominent changes in expression. In contrast, there was a remarkable increase in Cx36 immunoreactivity in *fmr1* mutants over this short developmental period (Figure 6E,H), which coincides with the emergence of spontaneous, electrically-coupled behavior. Specifically, we found a ∼45% increase in Cx36 puncta per *mnx1*:EGFP volume in *fmr1* mutant embryos compared to controls at 24 hpf. Taken with increased Ca^2+^ activity at this stage, elevated expression of gap junction proteins in *fmr1* embryos could mediate heightened electrical coupling and drive hyperactive motor output in this FXS model.

**Figure 6.**
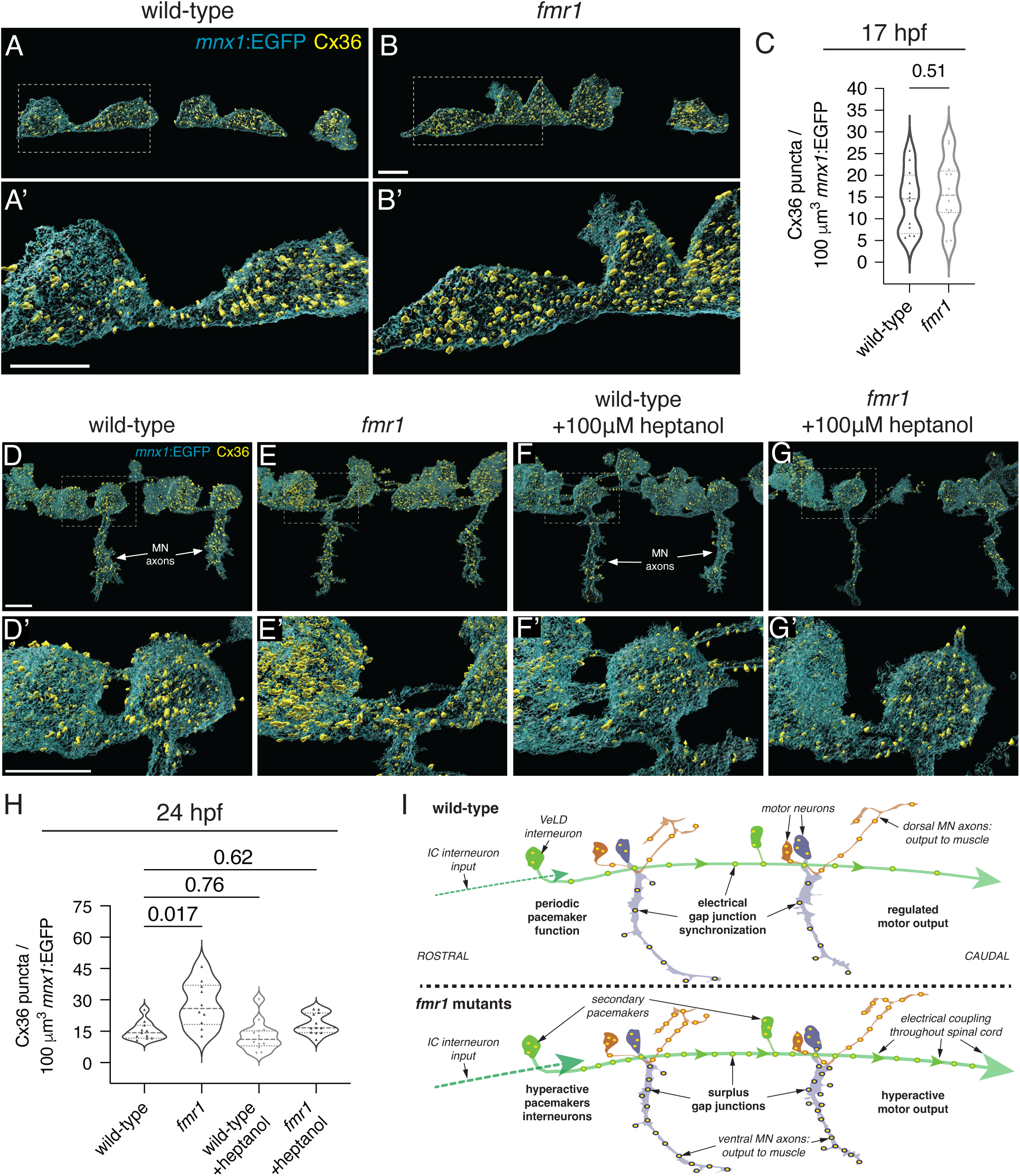
Increased expression of gap junction protein Connexin 36 in *fmr1* spinal motor circuit neurons is driven by electrical coupling. Images of transgenic *mnx1:*EGFP wild-type (A; A’ corresponds to zoom of dashed box in A) and *fmr1* mutant embryos (B,B’) processed to detect Cx36 protein at 17 hpf. Images display three-dimensional surface models of transgenic *mnx1*:EGFP+ cells (motor neurons and VeLD interneurons; cyan) and Cx36+ immunofluorescent puncta (yellow). (C) Quantification of average Cx36+ puncta per 100 μm3 of *mnx1:*EGFP volume at 17 hpf (wild-type=14.01±2.05 puncta; *fmr1*=16.05±2.2 puncta; significance determined by t-test; n=12 embryos each). Representative images of wild-type control (D,D’), *fmr1* control (E,E’) wild-type +heptanol (F,F’) and *fmr1* +heptanol (G,G’) *mnx1*:EGFP spinal motor circuits (cyan), with Cx36+ immunofluorescent puncta (yellow). (H) Quantification of Cx36 puncta per 100 μm3 *mnx1:*EGFP volume in control and heptanol-exposed conditions (wild-type control=15.20±1.31 puncta; *fmr1* control=27.79±3.41 puncta, wild-type +heptanol=12.51±1.94 puncta; *fmr1* +heptanol=18.38±1.43 puncta; significance determined by Kruskal-Wallis test with Dunn’s multiple comparisons test; nWT=11 embryos, n*fmr1*=10, nWT+Hep=13, n*fmr1*+Hep=12). (I) Model summarizing motor circuits in wild-type and *fmr1* mutant embryos (IC=ipsilateral caudal interneuron; VeLD=ventral lateral descending interneuron; MN=motor neuron). All surface models of Cx36 and *mnx1:*EGFP expression were generated and quantified with Imaris software. Cx36 puncta were located ≤0 μm to EGFP expression. Scale bars = 10 μm.

The *fmr1* gene encodes the RNA binding protein Fmrp, which is implicated in many activity-dependent neurodevelopmental processes^17,52,53^. The dramatic increase in Cx36 in *fmr1* mutants could suggest a repressive role for Fmrp in gap junction protein formation. To test a potential feedforward mechanism for electrical synaptic activity, we bath applied 100 μM heptanol on both wild-type and *fmr1* mutant embryos from 17-24 hpf and again quantified Cx36 expression on *mnx1:*EGFP^+^ cells. Although developmental heptanol exposure had no effect on Cx36 expression in wild-type motor circuits (Figure 6F,H), electrical blockade rescued Cx36 density in *fmr1* mutants to levels comparable to wild-type controls (Figure 6G,H). This may suggest that Fmrp represses gap junction expression in wild-type motor circuits, or that hyperactive motor circuits drive Cx36 expression. Human FMRP does not appear to bind *GJD2* RNA (which encodes Cx36)^54,55^, though the Cx36 antibody may detect an alternate gene product in our zebrafish experiments. Taken together, motor circuits in *fmr1* mutants are hyperactive, less sensitive to electrical coupling blockade, and have increased density of gap junctions (see Model, Figure 6I), thereby providing several potential foundations for hyperactivity in this FXS model.

## Discussion

The development of vertebrate motor circuits requires the synchronization of motor neuron activity across an extensive distance through the spinal cord to achieve coordinated locomotive output. Spontaneous motor contractions in zebrafish initiate through mechanisms that are intrinsic to the spinal cord^25,29^, and spontaneous activity influences motor axogenesis and synaptic maturation^7,56,57^. Pacemaker neurons (or central pattern generators) are the presumptive source of this spontaneous activity, generating rhythmic output in early neural networks to synchronize the firing patterns of immature neurons^10,26^. IC and VeLD INs are physiologically coupled with MNs and are presumed to drive the minimal motor circuit underlying early zebrafish behavior^25,26,35^, though these conclusions were largely rooted in physiological studies and the precise cellular relationships remained unclear. Our functional imaging studies solidify the IC pacemaker hypothesis in showing that persistent, rhythmic IC activity precedes Ca^2+^ events in the trunk spinal cord (Figure 2). In addition, the targeted removal of pioneer ICs lead to desynchronization of motor circuit activity, further demonstrating that ICs provide critical influence on developing circuits. VeLDs could be construed as secondary pacemakers, as they extend IC influence throughout the trunk spinal cord, showing tight synchronization with ipsilateral *mxn1^+^* MNs and caudal VeLDs as development progresses into widespread circuit activity (Figure 2). VeLD axogenesis appears to slightly precede IC axon outgrowth^32,58^, and VeLD physiology and morphology is remarkably similar to IC cells^25,26^. As early spontaneous contractions initiate at the tip of the embryo tail (see Videos 9, 10), IC interneurons - positioned in the caudal hindbrain - would require VeLDs to extend pacemaker influence through spinal cord. This interneuron network provides critical synchronization of early born MNs to kickstart locomotive activity.

We further uncovered important nuances in gap junction coupling as motor circuits coalesce, as we showed that subthreshold exposure to the gap junction inhibitor heptanol drove desynchronization of motor circuits and hyperactivity, in the form of uncoordinated motor output (Figs. 1,2). A previous study showed that concentrated heptanol exposure (2 mM) completely inhibits spontaneous coiling movements and blocks MN rhythmicity^8^, which suggests a complete saturation of gap junctions prevents pacemaker communication with MNs. In contrast, our subthreshold exposure paradigm showed that VeLD activity persists with partial electrical blockade, but these secondary pacemakers lose coupling with ipsilateral motor circuit cells (Figure 2). In line with this data, neuronal desynchronization is also noted in Cx36 and Cx40 mouse knockout models, which lack neuronal gap junction proteins^59,60^.

Importantly, our behavioral data also shows that partial gap junction blockade leads to weak, hyperactive output, akin to seizure-like behavior (Figure 1), which may present serious implications for gap junction-targeted therapeutics^61^. In line with this observation, retinal ganglion cells in Cx36 knockout mice display hyperactive low firing rates that contrast with concentrated, synchronized waveform activity in controls (though this phenotype did not influence eye-specific segregation)^62^. Our collective data suggests that rhythmic motor pacemaker output ensures coordinated locomotion through electrical synaptic connections, and partial disruption of gap junctions can lead to uncoordinated, hyperactive output.

What are the potential consequences for alterations in motor circuit formation? Some of the most debilitating symptoms of FXS and associated ASDs are repetitive motor behaviors and epilepsy^14,39,40^ and pathogenesis is likely rooted in neurodevelopmental mechanisms^63^. FXS model zebrafish display hyperactivity throughout embryonic, larval, and adult stages (Figure 4)^45,64^. We predict that hyperactivity in *fmr1* mutants is linked with excess gap junctions based on several findings. First, we found that subthreshold heptanol did not drive hyperactive behavior in *fmr1* mutants and did not influence Ca^2+^ event synchronization (Figure 5), effects that were prominent in wild-type (Figures 1,2). Second, we also showed increased gap junction Cx36 protein expression in motor circuit cells in *fmr1* (Figure 6). Interestingly, this increase in Cx36 in *fmr1* mutants appears to be driven by a feedforward mechanism, as heptanol blockade rescued Cx36 expression. It is unclear if this effect is directly influenced by the RNA binding protein Fmrp, absent in *fmr1* mutants^65^, though indirect effects are also possible through associated signaling cascades or epigenetics^66–68^. Although the mechanism remains unclear and there are many caveats to electrical synaptic therapeutics, restoration of gap junction expression in *fmr1* mutants via electrical blockade presents a powerful potential tool in the modulation of hyperactive motor circuits during development.

In the pathological context, there are a multitude of links between gap junctions and neurodevelopmental disorders. Elevations in astrocyte-associated gap junction protein CX43 are noted in autistic individuals^69^, and disruption of Connexin activity in human fetal tissue interrupts spontaneous cortical activity^70^. Connexins have also been suspected in the pathogenesis of epilepsy, with variations noted in gene and protein expression^71–73^. Zebrafish *connexin39.9* mutant embryos show reduced touch-evoked coiling behavior^51^, further indicating that electrical gap junction coupling drives early spontaneous behavior. However, hyperactivity in *fmr1* mutants could also be influenced by pacemaker activity, as *mnx1^+^* cells show increased Ca^2+^ activity and amplitude compared to controls (Figure 5). It remains to be determined if pacemaker hyperactivity lies directly upstream of increased Cx36 in *fmr1* mutants. Importantly, electrical coupling remains influential in more mature stages^48,49,74^, which suggests gap junctions could exert a persistent influence on hyperactivity in this FXS model. The emergence of chemical neurotransmission may also contribute to hyperactive locomotion phenotypes in FXS, as we’ve previously shown reduced expression of inhibitory synaptic machinery in *fmr1* embryos at the end of embryogenesis^50^. Therefore, inhibitory INs in our FXS model may have reduced influence as locomotive activity progresses into more complex swimming behavior. Our future work will center on the transition to chemical neurotransmission in motor circuits and test the degree that electrical coupling influences later stages of locomotion.

## RESOURCE AVAILABILITY

Additional information and resource requests will be fulfilled by the lead contact, Caleb Doll (<u>caleb.doll@</u> <u>cuanschutz.edu</u>). This study did not generate new unique reagents. Data reported will be shared upon request, along with any additional information required to reanalyze the data.

## EXPERIMENTAL MODEL

### For in vivo animal studies - zebrafish lines and husbandry

The Institutional Animal Care and Use Committee at the University of Colorado School of Medicine approved all animal work, which follows the US National Research Council’s Guide for the Care and Use of Laboratory Animals, the US Public Health Service’s Policy on Humane Care and Use of Laboratory Animals, and Guide for the Care and Use of Laboratory Animals. Larvae were raised at 28.5°C in embryo medium and staged as hours (hpf) according to morphological criteria ^28^. Zebrafish lines used in this study included *fmr1^hu^*^2787 75^, *Tg(mnx1:EGFP)^ml^*^2 37^, *Tg(mnx1:GAL4)^s^*^300t 76^, and *Tg(UAS:GCaMP-HS)^nk^*^4a 34^. Genotyping for *fmr1^hu^*^2787^ was performed as previously described ^44^. Only maternal-zygotic *fmr1* mutants were analyzed in our studies; due to the early birth date of VeLD and primary motor neurons^9^, this approach avoids the potential maternal contribution of *fmr1* RNA^77^. As zebrafish sex determination does not occur until juvenile stages ^78^, we were unable to determine the sex of the embryos in our experiments.

## METHOD DETAILS

### Drug treatments and behavioral analysis

At 24 hpf, embryos were removed from chorions and placed in petri dishes with egg water (control) or 1-heptanol dissolved in egg water (50 μM, 100 μM; Sigma-Aldrich). As embryos were dechorionated prior to cell ablations, 1-minute open field recordings were captured after 15 minutes of treatment using a SwiftCam SC1603 16-megapixel microscope camera (Swift Optical Instruments) attached to an iMac computer via C-Mount at ∼25 frames/second, in standard petri dishes. The water temperature and ambient temperature were recorded before, during and after each trial. The water temperature was maintained at 23±1°C whereas the ambient temperature was maintained at 24±1°C. Each recording was blindly analyzed frame by frame, and coiling behavior was scored based on the relative strength of coiling behavior, where “complete” coils represented a movement where the embryo tail reached the level of the hindbrain. In contrast, in “weak” coils an initiated coil did not result in the embryo tail reaching the level of the hindbrain (see Videos 1-3). Although heptanol can influence cardiac activity^79^, heart contractions in zebrafish embryos do not initiate until approximately 24 hpf and blood flow is not noted until around 36 hpf^80^.

### Calcium transient imaging and analysis

*mnx1:*Gal4;*UAS*:GCaMP-HS transgenic embryos were manually dechorionated at either 18-21 hpf or 23-26 hpf. Next, a razor blade was used to remove ∼5 mm of tissue from the caudal tip of the embryo tail. As the general anesthetic tricaine mesylate drives paralysis through voltage gated Na^+^ channels to reduce neuronal excitability^81^, we instead applied the curare-mimetic pancuronium bromide at 0.45 mg/ml concentration for 10 minutes, and then mounted embryos laterally in 1% low-melt agarose^82,83^. Changes in fluorescence in *GCaMP-HS*^+^ cells undergoing active Ca^2+^ events were captured at ∼65 ms resolution using a 40x water immersion objective and 126×126 pixel resolution for 1000 frames, on a Zeiss 880 microscope running Zen Black (Zeiss). Zeiss files (czi.) were converted to TIF format and the Matlab program EZcalcium was next used to correct motion drift in each video^84^. Next, the Matlab program FluoroSNNAP was used to generate unbiased representations of Ca^2+^ transients^85^. First, cell bodies of active cells were manually outlined to generate segmented images. Manual segmentation was required due to the sparsity of active cells, as the program was designed for more widespread activity. All videos were processed at 7 Hz frame rate (∼65 ms/framerate), using template-based event detection, a threshold of 0.85, and minimum ΔF/F amplitude of 0.01 (default settings). Baseline fluorescence was also defined with default settings (F_0_ at 10 seconds at the 50^th^ percentile). With these settings, the program automatically generated ΔF/F traces, amplitude, rise and fall times, and the number of Ca^2+^ events for each cell. Coupled and uncoupled peaks were quantified by comparing the relative Ca^2+^ event quantity for each cell in a circuit to the most active cell in that circuit, which were then weighted by the number of cells in the circuit. Example circuit:

> Cell 1 = 10 events; Cell 2 = 9 events, Cell 3 = 8 events
>
> Coupled events, 9+8 = 17; Uncoupled events, (10-9) + (10-8) = 3; Weight = 3 (total cells)
>
> Weighted coupled events = 17 / 3 = 5.67, Weighted uncoupled events = 3 / 3 = 1

Weighted values were summed for the entire dataset and significant changes in the relative proportion of coupled versus uncoupled peaks were determined using Fisher’s exact tests. Amplitude, circuit size, and events per cell were averaged by embryo and then compared using t-tests or Mann-Whitney tests based on the normality of the dataset.

### Cell ablations

*mnx1:*Gal4;*UAS*:GCaMP-HS transgenic embryos were manually dechorionated at the 17 hpf stage, then tail slits were performed prior to anesthesia with pancuronium bromide, as stated above. Embryos were then mounted in 1% low melt agarose with the dorsal spinal cord nearest the coverslip. Ablations were performed using a Zeiss 880 confocal microscope, using 405 and 488 nm lasers and a 40x objective, adapting an established protocol in zebrafish^38^. First, VeLD INs undergoing active Ca^2+^ events were located in the EGFP channel at 1x magnification. Next, digital zoom was increased to 30x and a circular region of interest (ROI) was placed on the active cell. Continuous scanning exposure of the ROI with 405 nm laser at 80% power for 45 seconds led to selective cell death. After ablation, digital zoom was reduced to 1x to ensure the absence of Ca^2+^ events following laser ablation. DIC imaging also showed cellular debris and nuclear fragmentation, further indicative of cell death. This was best exemplified in pilot experiments using a prenylated membrane reporter, which showed internalization of GFP expression alongside nuclear fragmentation (see Figure 3C-C’’)^38^. Embryos were then carefully removed from agarose and placed into prewarmed embryo media to continue developing until the ∼23 hpf stage (embryos harmed during agarose removal were discarded). At ∼23 hpf, tail slits were again necessary prior to pancuronium bromide anesthesia, and Ca^2+^ imaging was performed as described above.

### Embryonic behavior

At 20 somite-stage and 24 hpf, embryos within chorion membranes were distributed into 24 cells in prewarmed 1.5% agarose transplantation molds (PT-1, Adaptive Science Tools, Worcester, Massachusetts) filled with 500 μl of prewarmed egg water (28.5 C±0.5°C). The embryos equilibrated beneath the camera for 3 minutes before each recording. Video recordings were captured using a SwiftCam SC1603 16-megapixel microscope camera (Swift Optical Instruments, Schertz, Texas) connected via C-Mount adapter to a dissecting microscope. Images were acquired at ∼20 frames per second for 80 seconds. Temperature was maintained as described above. The video files were analyzed on a standard PC system with DanioScope software (version 1.0.109, Noldus, Leesburg, Virginia).

### Hatching

Embryos were collected and sorted soon after fertilization, placed in fresh egg water, and incubated at 28.5°C degrees. At 24 hpf, egg water was replaced with minimal disturbance to the embryos. At 48 hpf, hatching was quantified, as determined by the complete emergence of the embryo from its chorion. *Immunohistochemistry* Transgenic embryos expressing *mnx1:*EGFP were fixed in 4% paraformaldehyde/1xPBS and rocked overnight at 4°C. Embryos were washed twice for 15 minutes in 1xPBS/0.1%Triton X-100 (PBSTx), blocked 1 hour in 2%goat serum/2%bovine serum albumin/PBSTx, 200μL per slide. Samples were then incubated in rabbit α-Cx36 primary antibody (1:500, in block; 36-4600; Invitrogen), rocking overnight. Sections were then washed 6×15-minutes in PBSTx and incubated 2 hours rocking at room temperature in 200μL of Alexa Fluor goat α-rabbit 568 secondary antibody (1:500, in block; A-11011; Invitrogen). Embryos were washed for 1.5 hours (6×15-minute washes) in PBSTx, then mounted in 1% low-melt agarose for imaging.

### Immunohistochemical modeling and quantification

Immunohistochemical puncta were quantified from three-dimensional surface projections in Imaris x64 (v9.9.1; Oxford Instruments) from confocal z-stacks captured with identical settings: 40x water immersion Korr objective (NA = 1.20), 512 x 512 pixel frame, pixel dwell = 1.03 μs, 1.8 zoom, z-step = 0.208 μm, 488 laser power = 5.5%, 568 laser power = 6%, pinhole radius = 117 μm, using Airyscan Superresolution mode (Zeiss). In Imaris software, z-stacks were trimmed to include just one half of the spinal cord. Surfaces were built for *mnx1:*EGFP expression without smoothing, using an 8 μm background subtraction threshold. Next, the channel corresponding to Cx36 expression (568 nm) underwent deconvolution (standard parameters, robust iterative algorithm), for 10 iterations. Cx36 surfaces were then rendered without smoothing using thresholding with an average puncta diameter of 0.5 μm. Next, the background subtraction threshold was toggled by the user to reflect input fluorescence, and touching objects were split with an intensity-based method and seed point diameter of 0.5 μm. Surfaces were then generated using the maximum quantity of seed points and then filtered by size with a conservative lower filter of 0.25 μm^2^. Resultant surfaces were compared with original fluorescence to ensure proper fit. Next, Cx36 surfaces were filtered to retain only puncta ≤0 μm distant from *mnx1:*EGFP surfaces, to ensure proximity of gap junction proteins to motor circuit neurons. For each image, the total quantity of Cx36 puncta proximal to EGFP expression were recorded, which was normalized to the total volume of *mnx1*:EGFP expression in each sample and multiplied by 100 to provide average Cx36 puncta per 100 μm^3^ *mnx1:*EGFP volume.

## QUANTIFICATION AND STATISTICAL ANALYSIS

### Statistics

All statistics were performed in Graphpad Prism (version 9). Normality was assessed with a D’Agostino and Pearson omnibus test. For two groups, unpaired comparisons were made using either unpaired two-tailed t tests (for normal distributions) or Mann–Whitney tests (abnormal distributions). For more than two groups, comparisons were made with one-way ANOVA tests with Holm-Sidak’s multiple comparisons test (normal distributions), or a Kruskal-Wallis test with Dunn’s multiple comparisons test (abnormal distributions), with multiple comparisons made to wild-type controls. Sample sizes, raw data, and statistical details are available in figures or figure legends, where values = mean ± standard error. Violin plots display median and quartiles along with each datapoint.

### KEY RESOURCES

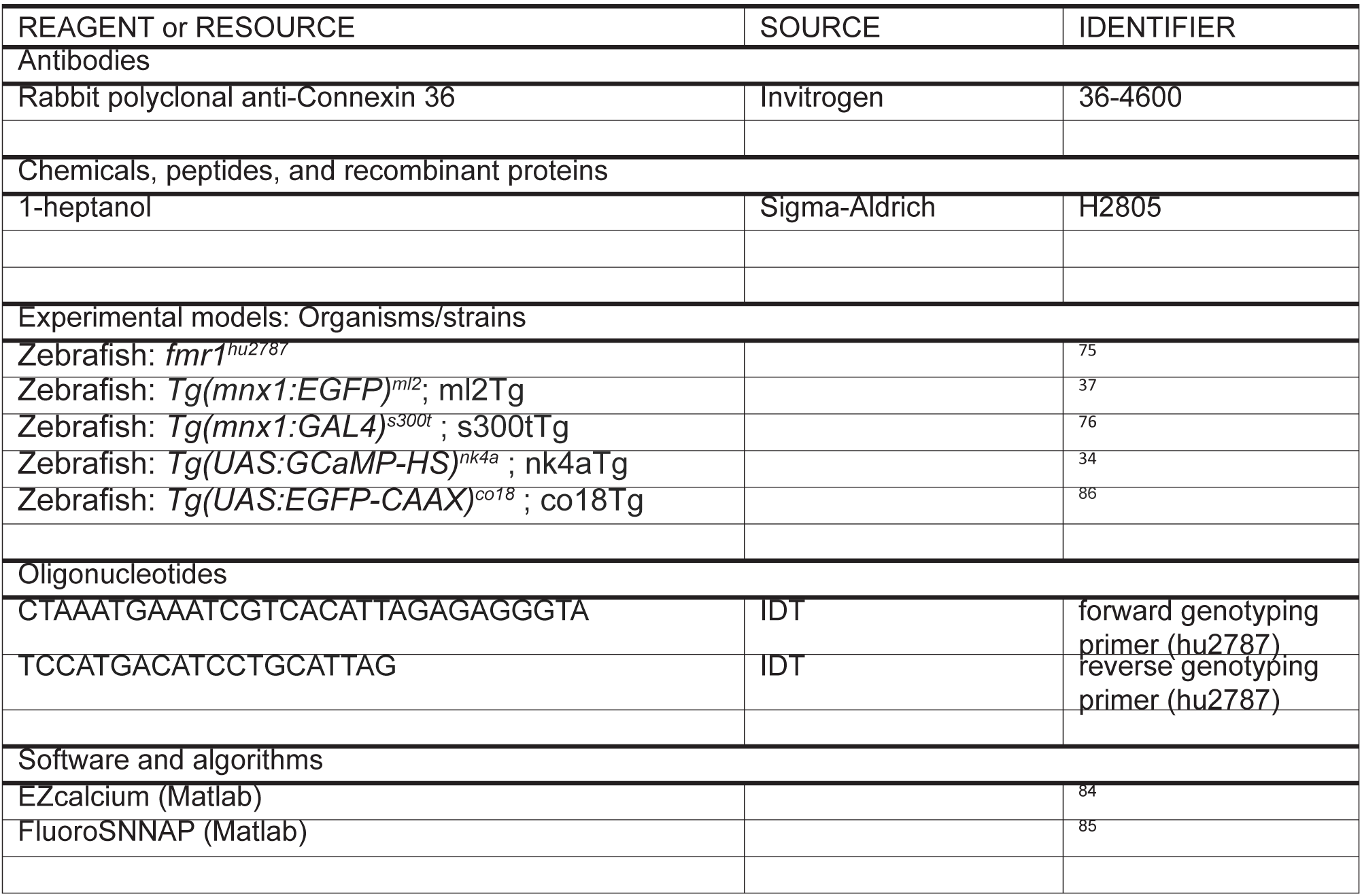

## Author contributions

KM: formal analysis and investigation. CB: formal analysis and investigation. BA: funding, editing. CD: conceptualization, formal analysis, investigation, writing, funding, visualization. All authors contributed to the article and approved the submitted version.

## Acknowledgements

We thank Natalie Carey and Oscar Mendez for insightful discussion; Angie Ribera, Melissa Wright, and Doug Hicks for transgenic lines; and the Appel laboratory for support and feedback.

